## Supplementary figuree for "Chickpea NCR13 disulfide cross-linking variants exhibit profound differences in antifungal activity and modes of action"

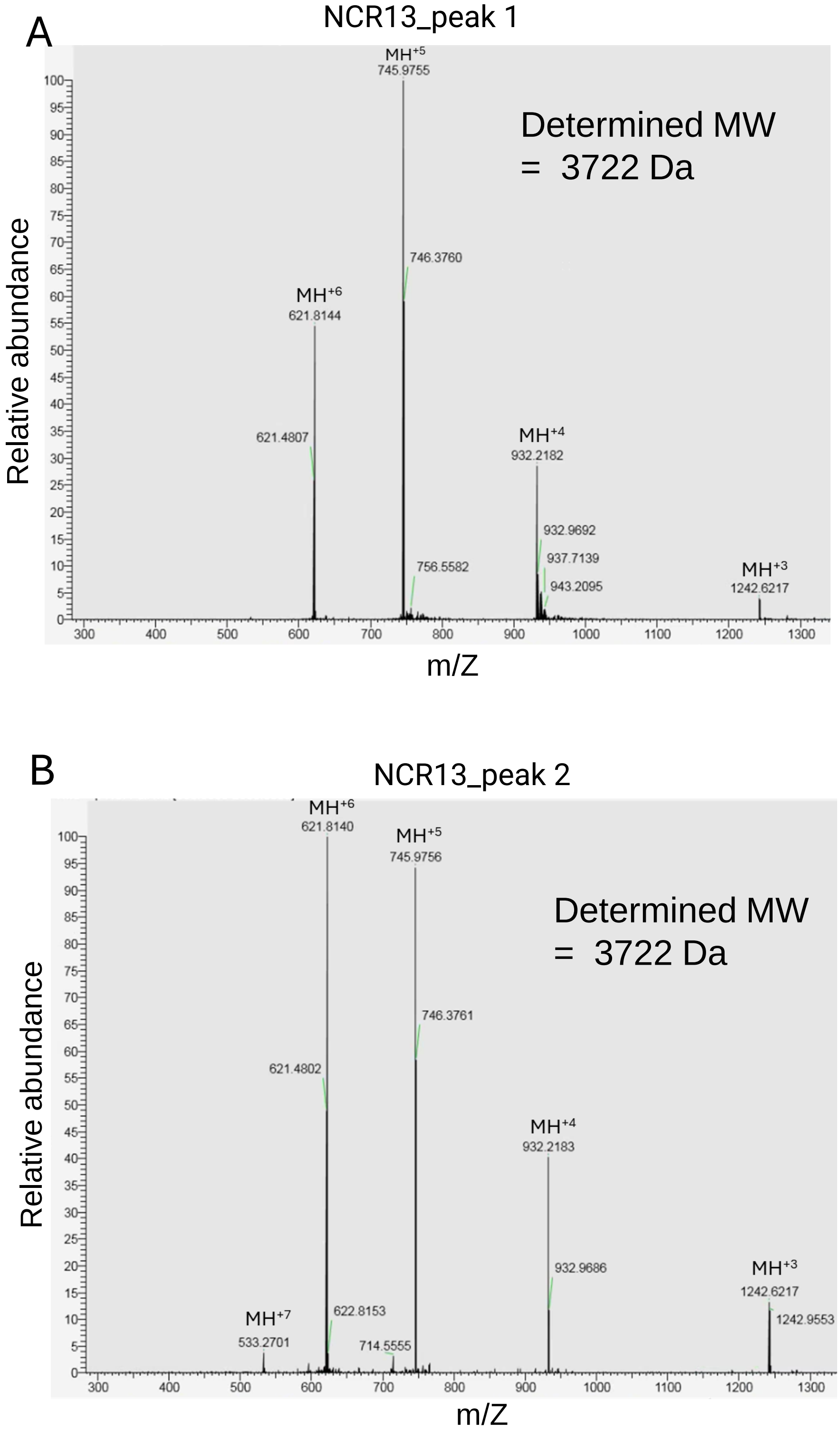


**Figure S1. Mass spectrometry confirms identical molecular weights for the products in NCR13_peak 1 and NCR13_peak 2.** (A-B) The mass spectra of NCR13_peak 1 and NCR13_peak 2 shows major peaks corresponding to different charge states of the peptide. MW = Molecular weight.

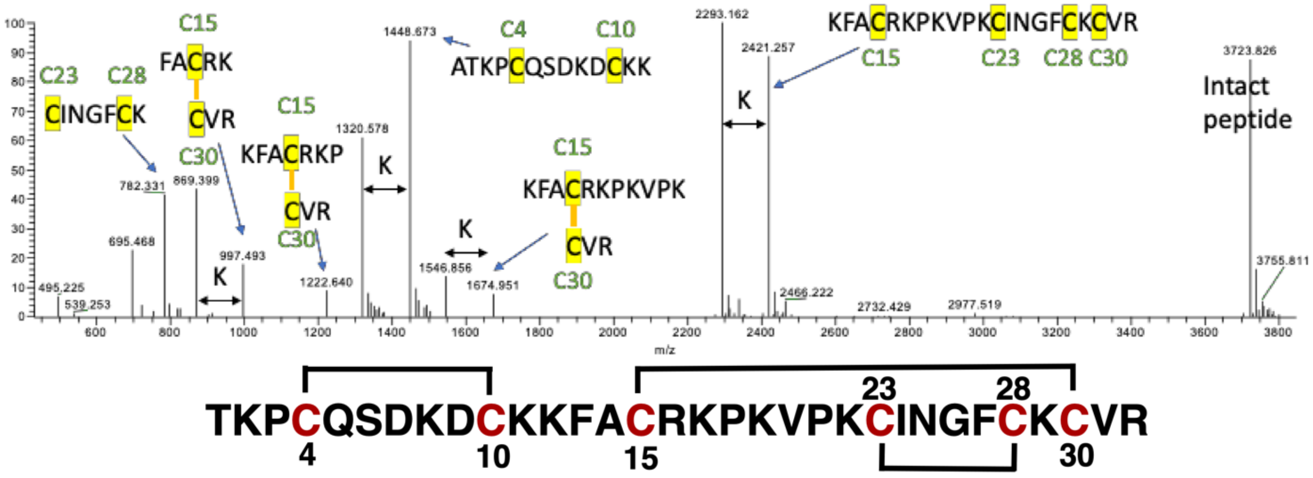


**Figure S2. Identification of the disulfide bond pattern for NCR13_PFV2 using mass spectrometry**. The primary amino sequence of NCR13 contains six cysteine residues which means there are 15 possible combinations of disulfide bond formation. In nature each NCRs are expected to fold with one specific disulfide bond combination. High-resolution mass spectrometry on a trypsin digested sample unambiguously show the pattern is C4-C10, C15-C30, and C23-C28 for NCR13_PFV2 as illustrated in the analysis above. There are at least three major digestion products that correspond to a C15-C30 disulfide and two corresponding to a C23-C28 disulfide. This information was crucial in solving the NMR solution structure for NCR13_PFV2.

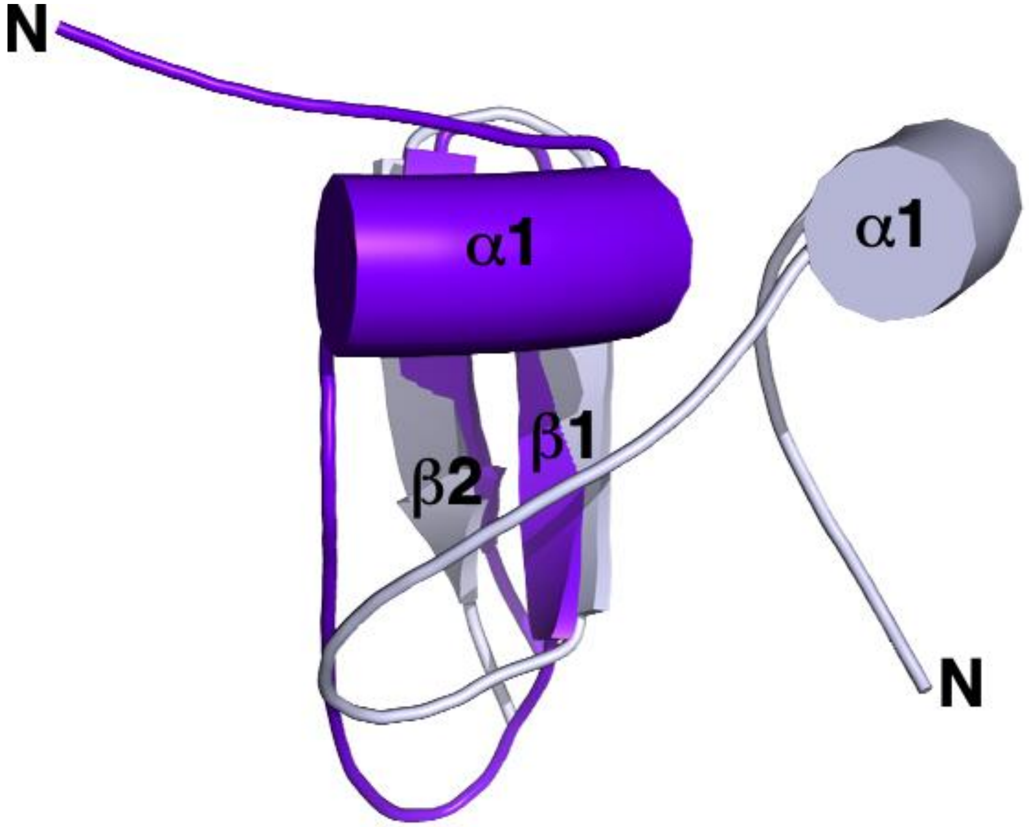


**Figure S3. NCR13_PFV1 and NCR13_PFV2 fold differently.** Cartoon representation of the structures of NCR13_PFV1 (purple) and NCR13_PFV2 (grey) superimposed on the β-sheet (V20 – V31). Helices are illustrated as cylinders. In NCR13_PFV2 the helix sits off to the side of the β-sheet while in NCR13_PFV1 it sits over the top of the β-sheet's face. This results in a slightly more compact structure for NCR13_PFV1.

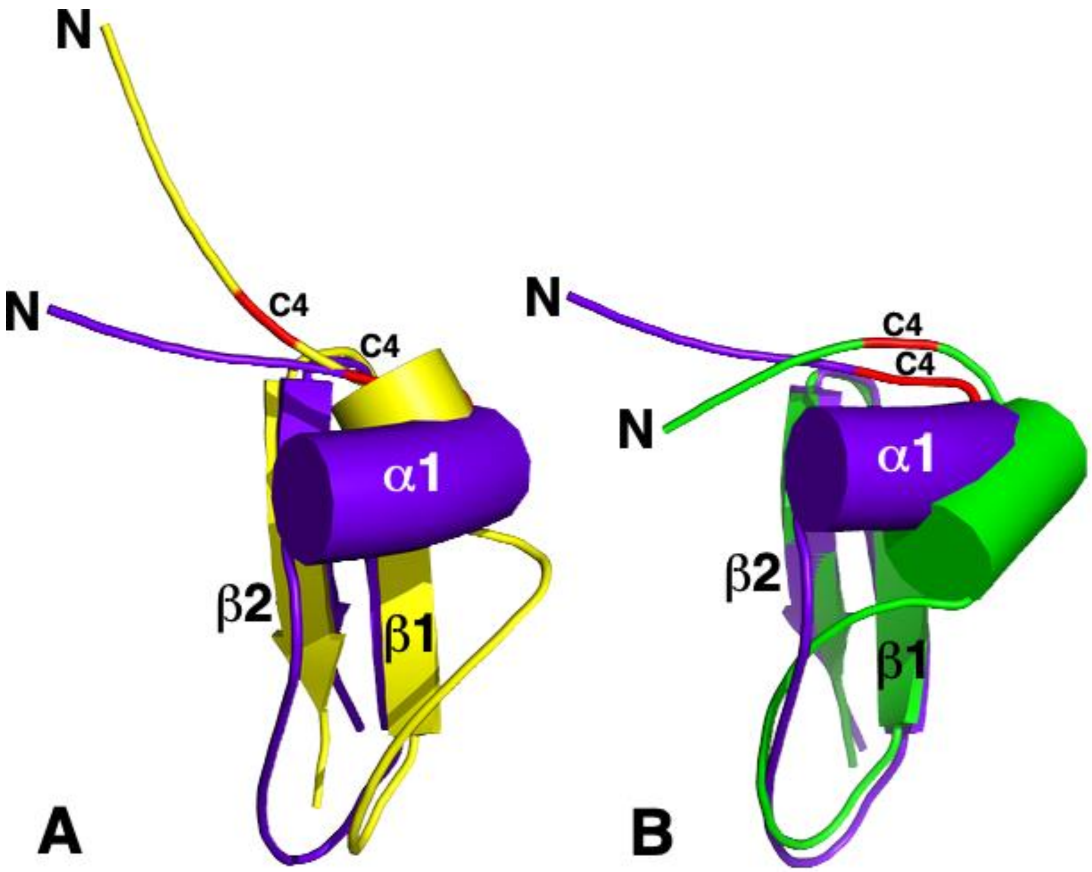


**Figure S4. Comparison of the experimental structure of NCR13_PFV1 with a structure calculated with a reversed disulfide bond pattern and with the predicted AlphaFold structure.** (A) Cartoon representation of the structures of NCR13_PFV1 (purple) and a NCR13_PFV1 structure calculated with a reverse disulfide bond pattern (yellow) superimposed on the β-sheet (V20 – V31). Helices are illustrated as cylinders with the position of C4 colored red in both structures. Given that the chemical shift data shows that both NCR13_PFV1 and NCR13_PFV2 contain a C15-C30 disulfide bond, there are two possible disulfide bond patterns possible for NCR13_PFV1: C4-C23/C10-28 (purple) and C4-C28/C10-C23 (yellow). Long range experimental NOEs between the N-terminal region of NCR13_PFV1 (T1-P3) and the β-strand were satisfied with the former disulfide pattern. In the reversed, C4-C28/C10-C23 disulfide pattern, the direction of α1 is also reversed relative to the β-strand and this prevents the N-terminal region (T1-P3) from making close contact with the β-strand. (B) Cartoon representation of the structures of NCR13_PFV1 (purple) and the AlphFold predicted structure (green) superimposed on the β-sheet (V20 – V31). Helices are illustrated as cylinders with the position of C4 colored red in both structures. While the β-sheet is similar in both the experimental and predicted structures, AlphaFold predicted a slightly shorter α-helix displaced two residues (D7-K11 versus Q5-C10 experimentally) that packs against the β-strand a bit differently. Note that the N-terminal region (T1-P3) folds towards the β-strand in the AlphaFold predicted structure as observed experimentally.

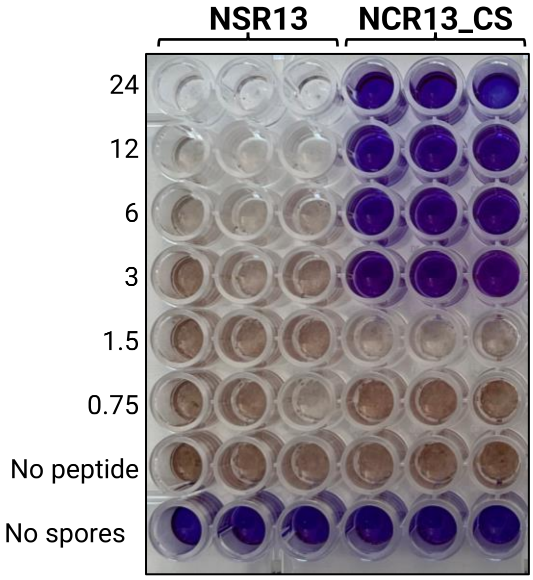


**Figure S5. Removing all six cysteine residues in NCR13 abolishes activity against *B. cinerea***. Comparison of the antifungal activity of a full disulfide knockout, NSR13, and chemically synthesized NCR13 (NCR13_CS) Fungal cell viability assay performed with resazurin. A change from blue to pink/colorless signals resazurin reduction and indicates metabolically active *B. cinerea* germlings after 60 h.

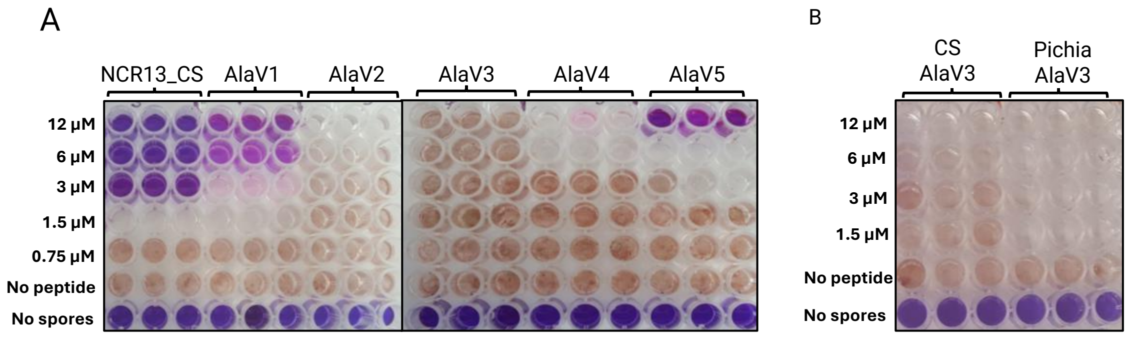


**Figure S6. Identification of the NCR13 active motif.** (A) Antifungal activity of synthetic NCR13 and NCR13 alanine mutant variants. The latter constructs were generated by substituting a window of alanine residues within the NCR13 core sequence, leading to the creation of constructs NCR13_AlaV1 through V5. (B) Comparison of the antifungal activity of chemically synthesized NCR13_AlaV3 *and P. pastoris* produced NCR13_AlaV3. All experiments were performed against *B. cinerea* using the resazurin fungal cell viability assay. A color change from blue to pink/colorless signals resazurin reduction indicating metabolically active fungal spores after 48 h. For each concentration of peptides, three biological replicates were used. Calculated MIC values are provided on the side of each experiment.

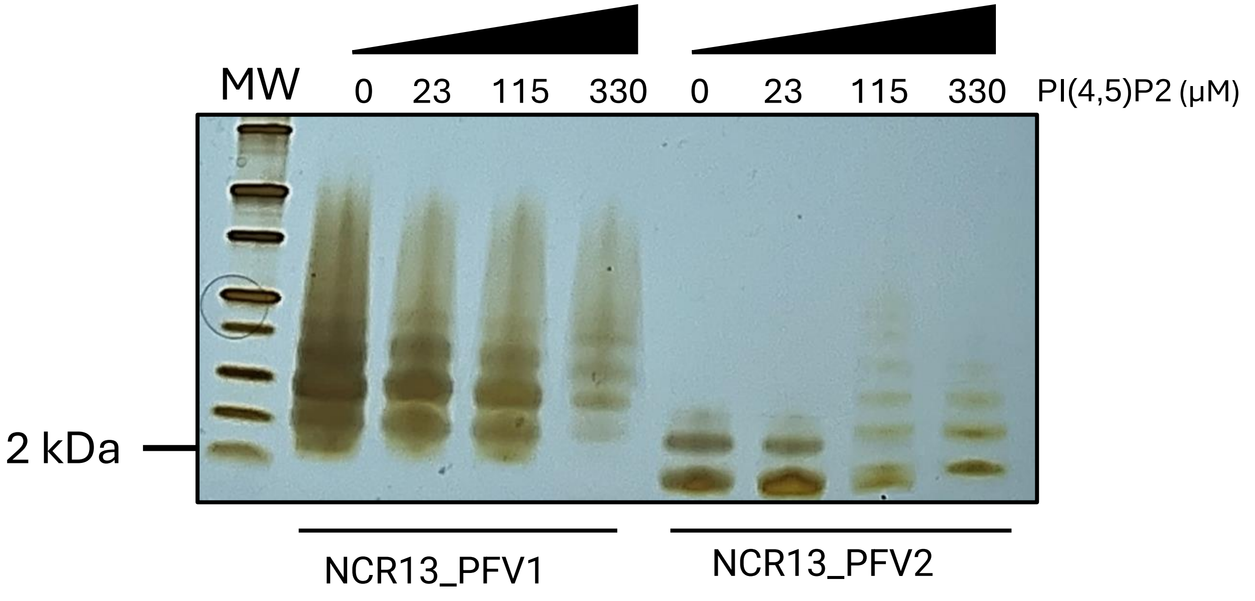


**Figure S7. Peptide oligomerization in the presence of PI(4,5)P2.** To determine if the peptide oligomerized in the presence of lipid, chemical crosslinking experiments were performed with NCR13_PFV1 and NCR13_PFV2 in the presence of different concentrations of PI(4,5)P2. Followed incubation with the biochemical cross-linker bis(sulfosuccinimidyl)suberate (BS3), the peptide were run on an SDS-PAGE gel and the product visualized by silver staining. MW indicates molecular weight protein marker. Shown image is representative of three independent experiments.

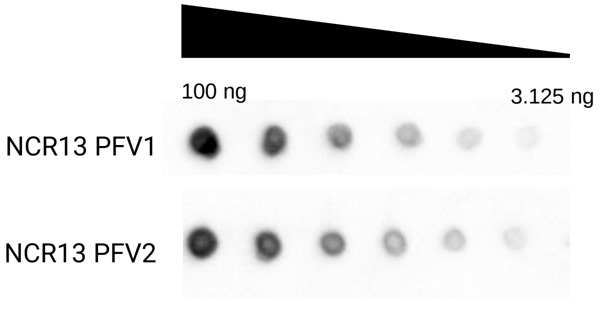


**Figure S8. NCR13 antibodies exhibit similar binding affinity to both disufide variants, PFV1 and PFV2.** Dot blot analysis of purified NCR13_PFV1 and PFV2 using anti-NCR13 antibodies (0.1ug/mL) followed by goat anti-rabbit IgG HRP (Cytiva RPN4301) at 1:20,000 dilution.

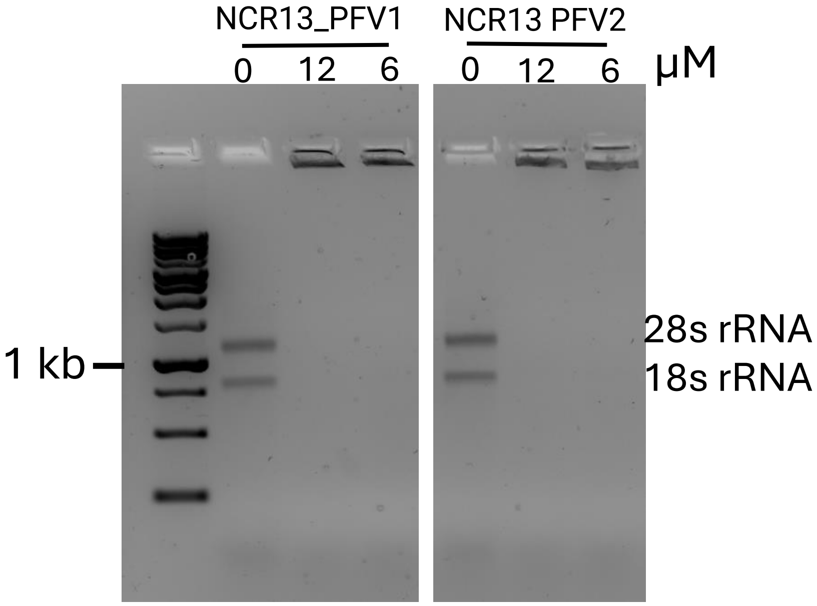


**Figure S9. NCR13_PFV2 exhibits rRNA binding at higher concentrations.** Electrophoretic mobility shift assay (EMSA) to determine if NCR13_PFV1 and NCR13_PFV2 bind rRNA. *B. cinerea* 28s and 18s rRNA was used to assess binding by electrophoresis on an agarose gel (1%). Peptide concentrations are indicated above the lanes. The first lane on the left contains molecular weight markers.

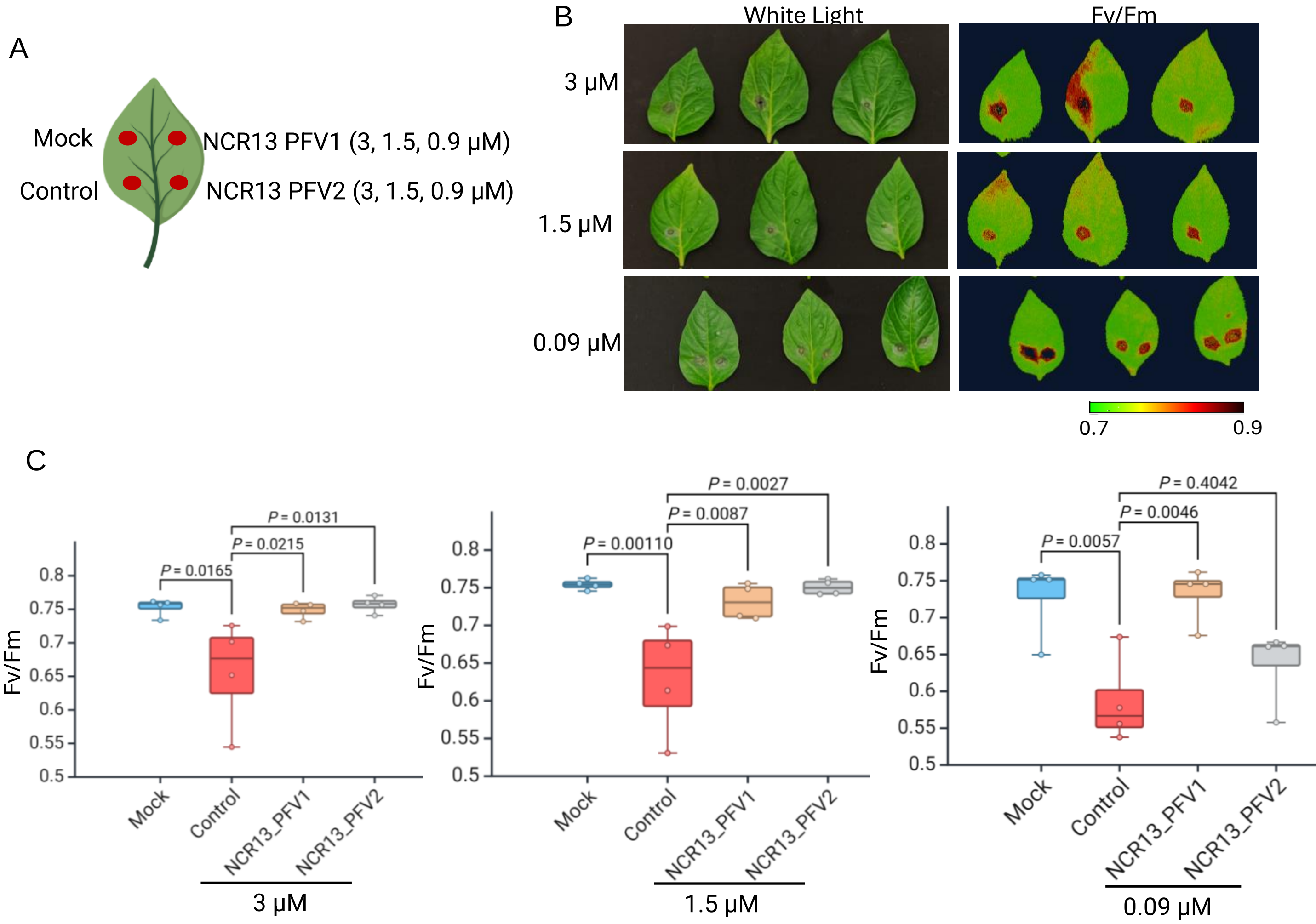


**Figure S10. Semi-*in planta* antifungal activity of NCR13_PFV1 and NCR13_PFV2 against *B. cinerea* on detached pepper leaves** (A) Model of each treatment location on the pepper leaf (B) Representative pictures (under white light and with CropReporter) showing the antifungal activity of NCR13_PFV1 and NCR13_PFV2 at 3, 1.5, and 0.09 µM against *B. cinerea* on detached pepper leaves. N = 4, N refers to biological replicates. (C) Photosynthetic efficiency (Fv/Fm) measurements of diseased lesions. In the box plot, Horizontal lines represent the median and boxes indicate the 25th and 75th percentiles. Statistical significance between control and treated samples were tested using One way ANOVA with Dunnett Multiple comparison test.

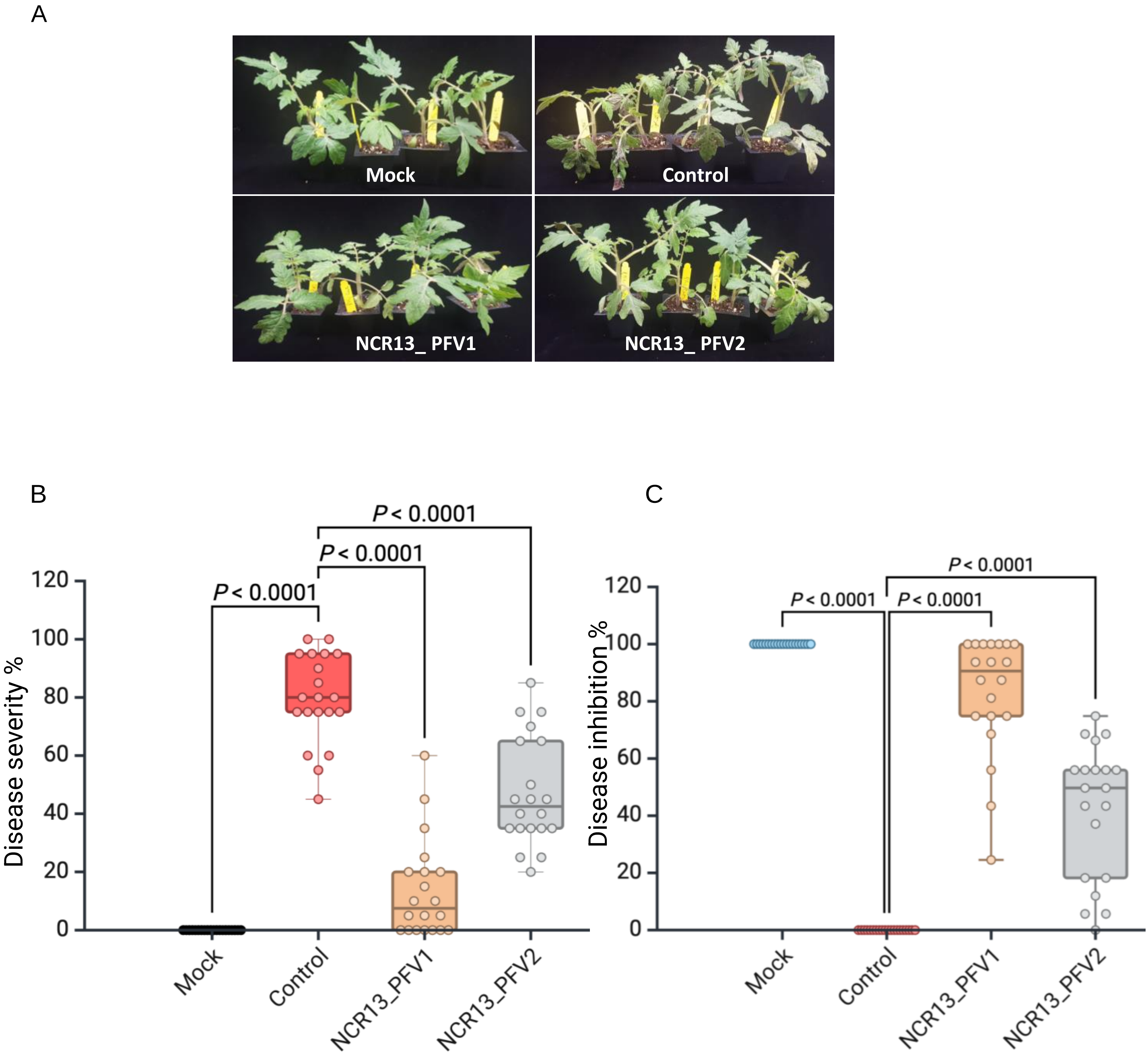


**Figure S11. *In planta* curative antifungal activity of NCR13_PFV1 and NCR13_PFV2 against *B. cinerea* on tomato plants.** Four-weeks old tomato plants sprayed with 1 mL of a 5 × $10^4$ *B. cinerea* spores suspension followed 24 hours later by a spray containing either 2 mL of water or NCR13_PV1 or NCR13_PV2 at 3 µM. (A) Representative pictures showing the curative antifungal activity of NCR13_PFV1 and NCR13_PFV2 at 3 µM against *B. cinerea* on tomato leaves. (B) Disease severity %; (C) Disease inhibition %. For panel (C) and (D) each data point represent mean ± SEM denoted by a dot. the average of 5 leaves per plant for 4 plants per treatment. Statistical significance between control and treated samples were tested using One way ANOVA with Dunnett Multiple comparison test. Three independent experiments were conducted with similar results.
